# An Adhesive-Based Fabrication Technique for Culture of Lung Airway Epithelial Cells with Applications in Microfluidics and Lung-on-a-Chip

**DOI:** 10.1101/2020.11.19.390674

**Authors:** Nicholas Tiessen, Mohammadhossein Dabaghi, Quynh Cao, Abiram Chandiramohan, P. Ravi Selvaganapathy, Jeremy A. Hirota

**Affiliations:** McMaster University; McMaster University/Infinotype

**Author notes:** Both authors contribute equally to this work (Nicholas Tiessen, Mohammadhossein Dabaghi).

## Abstract

This work describes a versatile and cost-effective cell culture method for growing adherent cells on a porous membrane using pressure-sensitive double-sided adhesives. This technique allows cell culture using conventional methods and easy transfer to microfluidic chip devices. To support the viability of our system, we evaluate the toxicity effect of four different adhesives on two distinct airway epithelial cell lines and show functional applications for microfluidic cell culture chip fabrication. We showed that cells could be grown and expanded on a “floating” membrane, which can be transferred upon cell confluency to a microfluidic chip for further analysis. The viability of cells and their inflammatory responses to IL-1β stimulation was investigated. Such a technique would be useful to culture cells in a conventional fashion, which is more convenient and faster, and stimulate cells in an advanced model with perfusion when needed.

## 2 Introduction

Chronic respiratory diseases have significant impacts on the health of the global population. Among the most significant, chronic obstructive pulmonary disease (COPD) is the third-highest cause of mortality internationally, accounting for over 3 million deaths annually[1]. Furthermore, greater than 300 million individuals are asthmatic, and 1 in 25,000 Caucasian individuals are born with Cystic Fibrosis (CF)[1]–[4]. In the lung, the airway epithelium provides the first line of defense, and dysfunction of this tissue is directly associated with the aforementioned chronic respiratory diseases[5]. The airway epithelium provides protection through many facets, such as mucociliary escalation, paracellular permeability regulation, and innate immunity[5], [6]. The airway epithelium is increasingly a target for new treatment options for the management of respiratory diseases. As a result, airway epithelial cell culture systems are poised to remain essential tools in basic research and drug discovery.

Conventional cell culture systems for adherent cells are based on growing cells on flat surfaces, including well-plates, culture dishes, microscope slides, and culture flasks[7]. Although these platforms have been widely applied, they have low flexibility in design and architecture[8]. In addition, epithelial cell biology has mostly been studied using static culture where cells exist in either submerged monolayer or air-liquid interface systems with media feeding from the apical or basal cell surfaces, respectively. Both of these models are suitable for basic and translational research and readily accessible. However, in their most common incarnations, these models do not integrate dynamic forces associated with airflow or interstitial fluid flow that are observed *in vivo*, which is likely a crucial component that impacts our basic understanding of airway epithelial cell biology in health and disease [9]–[12].

To improve existing cell culture model designs, advancements have been made in the miniaturization of cell culture models[13]–[17]. Miniaturized cell culture models provide suitable microenvironments to study live cells with a significant reduction in reagents and improved ability for high-throughput and control of model design[7]. Another progression in cell culture models has been the creation of miniaturized organ-on-a-chip devices[16]. These microfluidic culture systems integrate physiological forces of flow and mechanical strain that provide an interpretation of *in vivo* conditions[9], [11], [16]. In addition, the models have been advanced to incorporate co-culture platforms that simulate the microenvironments of cell-cell interactions[9]. Responses of airway epithelial cells have been different in such dynamic models when comparing to conventional culture systems[9], [11], [12]. Thus, the development of advanced models that integrate physiological forces for interrogating biological processes may have significant impacts on our basic understanding of cell biology through to drug discovery. Current organ-on-chip models frequently require cells to be seeded into a fully fabricated device, which requires new cell culture techniques rather than allowing the user to utilize standard cell culture methods[9], [10], [16], [18]. In addition, seeding cells into a fabricated model restricts the scalability of the system and the ability to access the cells after perfusion. Therefore, limitations currently still exist that need to be addressed to expand the applicability and uptake of these model systems.

While reported perfusion models do provide a platform to study cells with perfusion and integrate relevant forces on the cells, the materials and methods required to fabricate them limit their uptake into biomedical research labs[10], [16], [17], [19]. Exploring novel methods to assemble perfusion devices could have significant implications for expanding research possibilities in HAEC studies. Adhesive tapes come in a wide variety of chemical compositions and have varying physical characteristics[20]. Exploring the use of adhesives as a method for developing a cell patterning platform and a method for adhering layers of a perfusion chip may provide a possible new method that is accessible and addresses the mentioned limitations of methods that are currently available for HAEC culture in these spaces. To develop an airway epithelial cell culture model capable of organ on chip designs, our group has investigated the properties of four commercially available adhesives in varied cell culture conditions. We demonstrate that adhesives can be used as a tool for the modular assembly of perfusion cell culture devices without prior cell seeding. The adhesive model design presented is amenable to microscopy during experimentation and is easily deconstructed to allow access to cells post perfusion.

## 3 Results

### 3.1 Adhesives do not impact airway epithelial cell viability

The tested adhesives were characterized for toxicity to determine if the chemical composition of the adhesive material had negative effects on cell viability when they were submerged in culture media with HAECs. A decrease in green fluorescence would indicate a decrease in viable live cells when stained with Calcein AM. Qualitative visual analysis of Calu-3 cell and HBEC6-KT viability with four commercially available adhesives did not display any significant changes in viability, as shown with Calcein AM viability dye (Figure 1A). Although dark regions are shown in some panels when the cells were stained with Calcein AM, these changes are attributed to the 3-dimensional cell culture behavior of HAECs when they reach confluency. In addition, the commonly described “cobblestone” morphology of the HAECs was maintained in all conditions (Figure 1A). Furthermore, the qualitative analysis of total viable cells grown with the four adhesives did not yield any significant difference between the adhesive-grown conditions in comparison to a control (Figure 1B and C). Lastly, LDH assay results confirmed that adhesives did not compromise cell viability (Figure 1D and E). Therefore, the four commercially available adhesives did not show any negative effects on HAEC viability when grown in submerged culture conditions.

**Figure 1:**
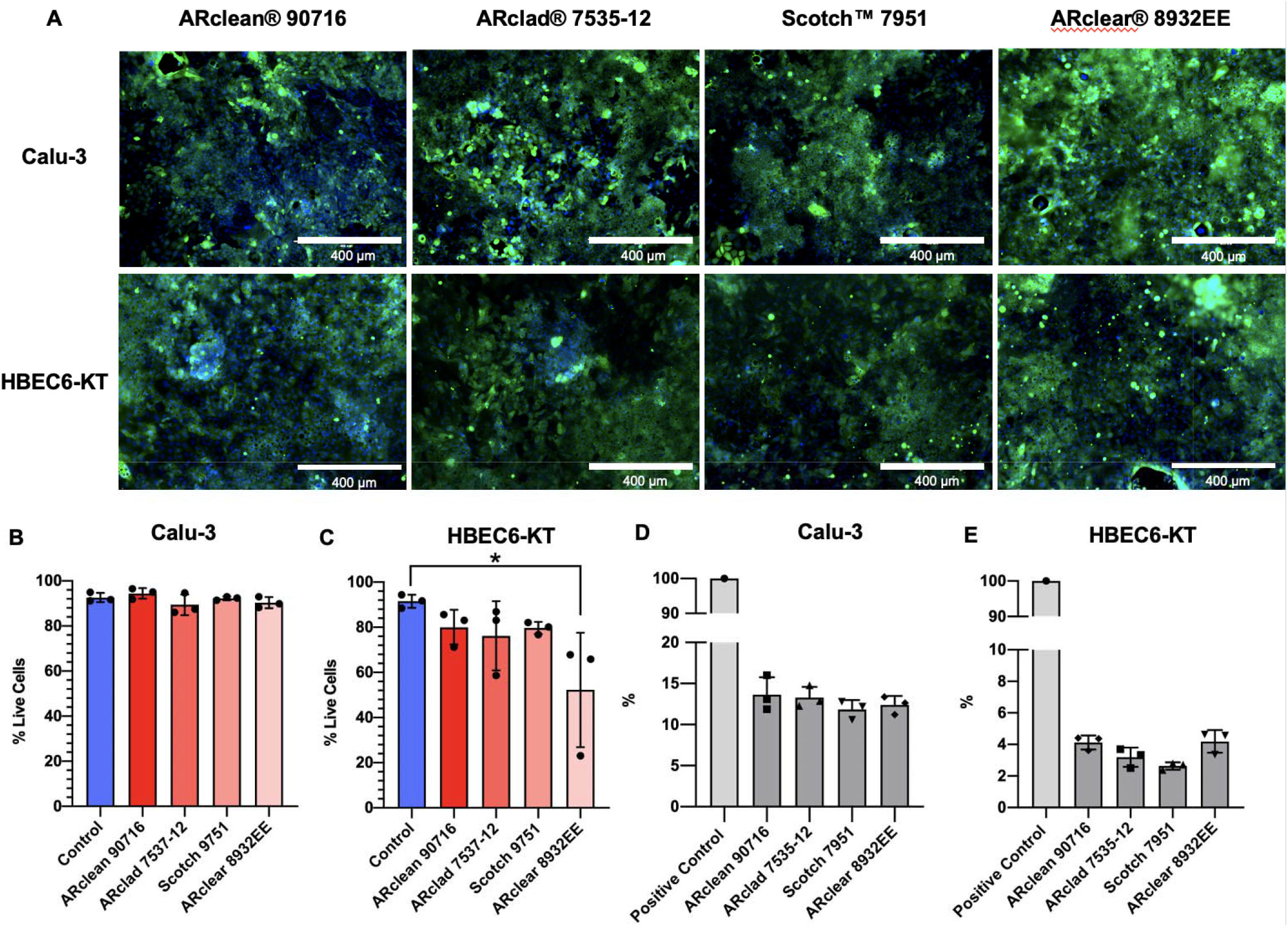
Qualitative characterization of cytotoxicity of several adhesives using both Calu-3 and HBEC6-KT cells. (A) Calu-3 and HBEC6-KT cells grown on top of ARclad® 7535-12, ARclean® 90716, Scotch™ 7951, or ARclear® 8932EE adhesives, respectively. The following growth to confluency, the cells were incubated in PBS with Calcein AM viability dye (green) and Hoechst nuclear dye (blue) at 37°C for 20 minutes. Images were taken at 100X magnification, and scale bars in the bottom right corner represent 400 μm. (B and C) Percent of viable cells following adhesive exposure, as determined by trypan blue live/dead stain. (D and E) LDH assay after growth for 7 days adhesives expressed as a percentage of a lysed positive control (n=3).

### 3.2 Microfluidic Device Fabrication

The microfluidic model consists of two layers of adhesive, a cell growth membrane, and a molded lid made of PDMS adhered together in a sandwich-like manner to a glass slide that provides optical clarity for microscopy and structural support of the chip as shown in Figure 2A. The mold for the PDMS lid was fabricated by conventional photolithography using a cleanroom facility. PDMS was prepared and cured on the top of the mold according to the manufacturer’s directions. Before pouring PDMS into the mold, two pieces of silicone tubing were placed on the designated locations as each chip inlet and outlet. This technique was applied for all functional experiments using adhesive assembled chips. The perfusion channel was designed with a 1mm channel that opens into a 5mm diameter circular growth chamber. The depth of the channel was 150μm. For all of the functional experiments, a 5 mm diameter circular growth area was cut out of the top adhesive. The workflow for adhesive chip assembly involves several steps, as presented in Figure 2B. The process begins with cutting a 0.5cm diameter circle into ARclean® 90716 adhesive on a Silhouette CAMEO® 3 (Figure 2B1). Both seals are removed from the bottom adhesive, and only the bottom seal is removed from the top adhesive. A porous polyester membrane is adhered between the two layers of adhesive and compressed with a hand roller (Figure 2B2). Following cell growth to confluency (Figure 2B3), the top seal of the top adhesive was removed, and the molded PDMS lid is attached to the chip (Figure 2B4). Then, the assembled chip with cultured cells was ready to be perfused (Figure 2B5).

**Figure 2:**
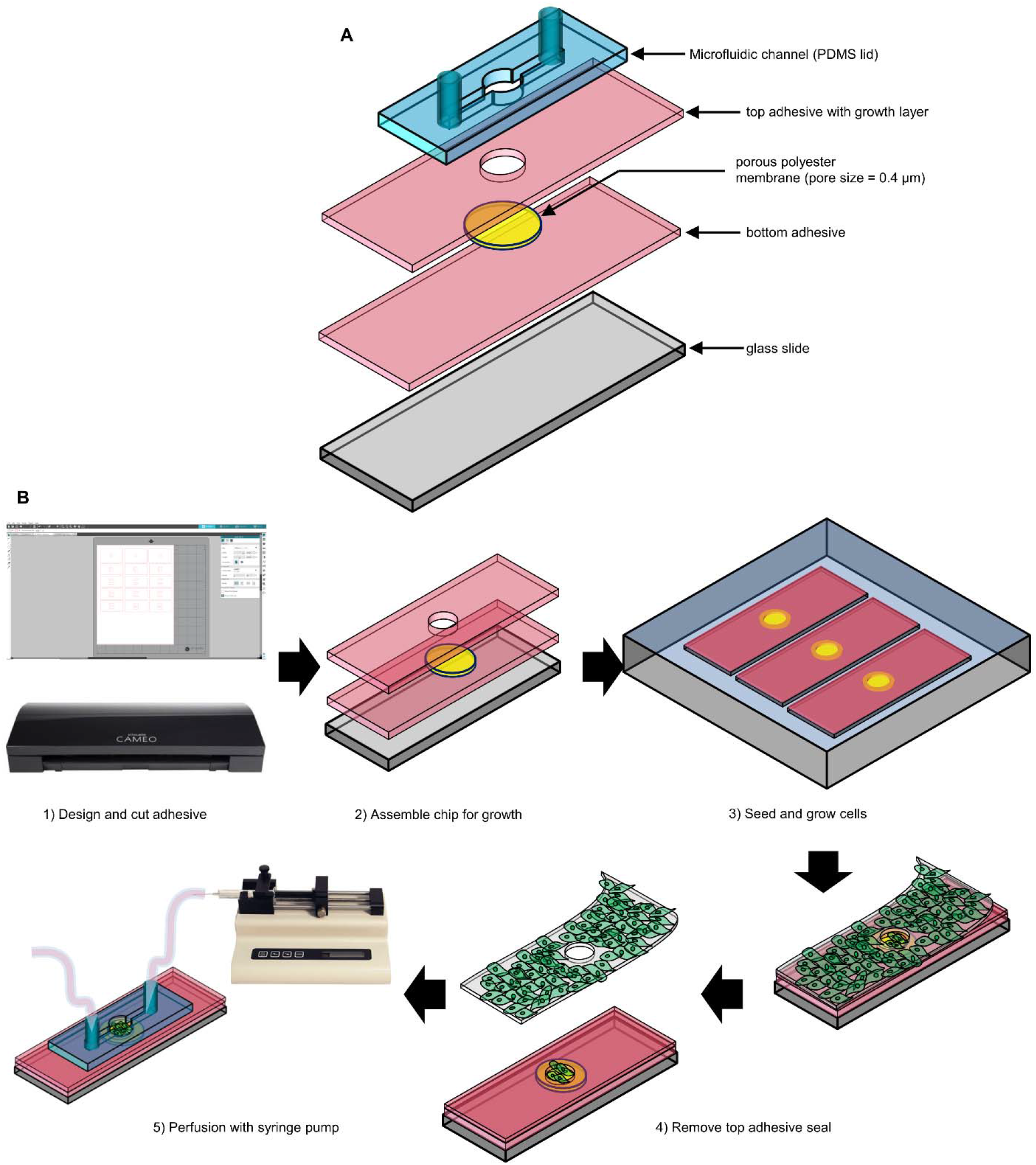
Diagram of the adhesive based epithelial cell chip manufacture process and workflow. (A) Explosion diagram of the stacked adhesive chip, including all layers. (B) Basic design methods and workflow for assembling and growing epithelial cells in the adhesive based chip.

### 3.3 Adhesive assembled microfluidic devices offer robust physical characteristics

Since the adhesive with growing cells was kept under a wet condition (submerged in a media to maintain the cells) and they were still wet (even after peeling off the liners) upon attachment to the PDMS lid, it was essential to evaluate the quality of this “wet” bonding between the wet adhesive and the PDMS lid. First, we conducted a flow experiment to ensure that there was no flow leakage.

The proposed adhesive assembled microfluidic model remained structurally viable after 30 minutes of several flow rates (10, 50, and 250μL/min) after being submerged for one week, simulating cell culture conditions (Figure 3A). After 30-minutes of water perfusion to the chips attached to the wet adhesives, the collected water at the outlet was weighted, and the flow rates were calculated, showing no noticeable difference between the measured flow rates and the set flow rates.

**Figure 3:**
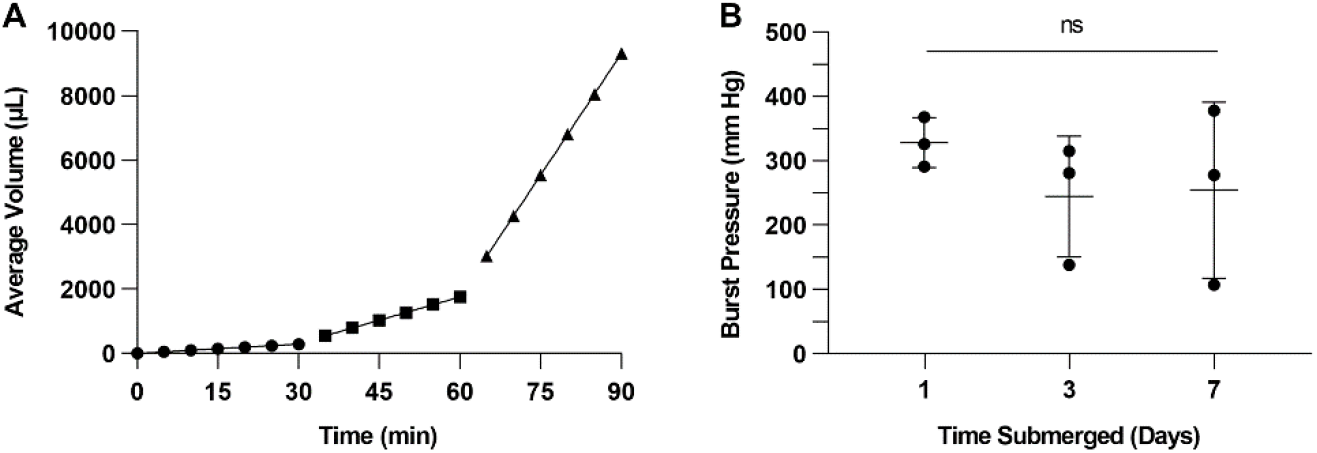
Volumetric flow validation and burst pressure analysis of ARclean® 90716 assembled perfusion chip. (A) A collected volume of water flowed through adhesive assembled chips submerged in α MEM for one week at 37°C. Water was flowed through the chip with a syringe pump for 30 minutes at 10 (circles), 50 (squares), and 250 (triangles) μL/min. Lines represent segmented linear regression of each flow rate. Bars for SD were plotted but were too small to be shown (n=3). (B) Burst pressure of adhesive assembled chips following submersion in growth media for 1-7 days. Burst pressure is shown for all replicates of the three conditions, with bars showing SD and mean of the groups (n=3).

Next, the burst pressure of the assembled chips that were fabricated using a similar method was measured. The model withstood burst pressures of 328.33 ± 31.48, 244.67 ± 76.69, and 254.33 ± 111.89 mm Hg for 1, 3, and 7 days of submersion, respectively (± SD, n=3, Figure 3B). These results suggested the wet bonding would be strong enough to achieve a leak-free perfusion.

To utilize the perfusion model for cell culture purposes, the maintenance of cells in a targeted growth area was desired to prevent cells from decreasing adherence of the PDMS lid to the top adhesive layer. Our proposed method of restricting Calu-3 cells in a targeted growth area (5mm diameter circle) was consistent in maintaining a defined interface of cell presence within the target area after adhesive seal removal (Figure S1 in the supplementary).

Calu-3 cells were qualitatively demonstrated to remain viable, as shown with Calcein AM viability dye, in the adhesive assembled device after perfusion of media for 24 hours (Figure 4). The post-perfusion images were taken by removing the lid, demonstrating an application of the model to access viable cells after sustained periods of perfused media.

**Figure 4:**
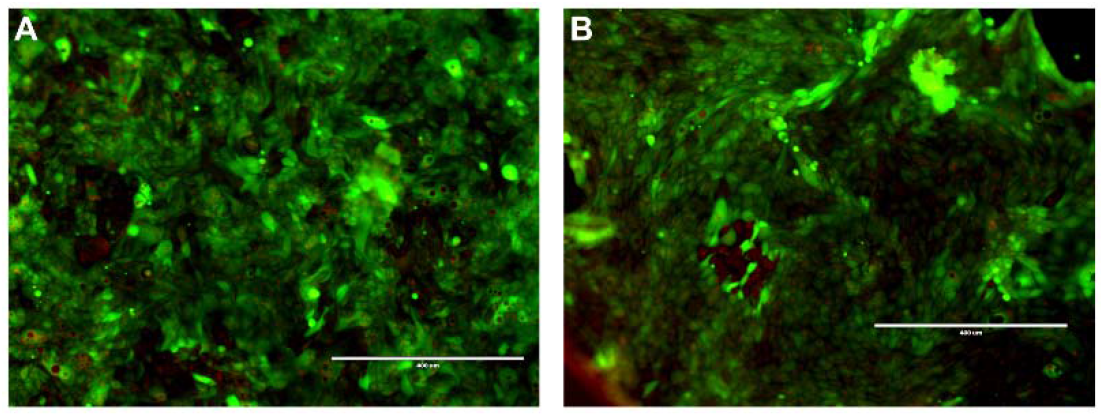
Qualitative Calu-3 cell viability following growth with media perfusion in an ARclean® 90716 adhesive assembled chip. (A) Cells were grown to confluency prior to perfusion of media and stained with Calcein AM viability dye (green) and NucRed™ Live 647 ReadyProbes™ Reagent (red) in PBS. (B) Cells following 24 hours of media perfusion at a flow rate of 20μL/min, stained as previously described. Images were taken at 100X magnification, and scale bars represent 400 μm.

Calu-3 cells were used to determine if a known inflammatory cytokine release after stimulation with IL-1β (5ng/mL) would elicit different responses in a perfusion culture system when compared to controls. In the perfusion culture system, IL-1β stimulation produced a 9-fold increase of IL-8 compared to control perfusion (p<0.01), as depicted in Figure 6A. In the perfusion IL-1β stimulation condition, IL-6 release increased significantly compared to their respective controls (p<0.0001)(Figure 6B).

### 3.4 Fabrication of lung-on-a-chip devices using Adhesive technique

To challenge and advance the wet boding, Calu-3 cells were cultured and expanded on porous polyester membranes in a static cell culture platform, followed by transferring these membranes to microfluidic chips. First, one mL of the cell culture media containing Calu-3 cells with a density of 5×10^5^ cells mL^−1^ was dispersed on a porous polyester membrane (membranes had a diameter of 25 mm and they were placed in a 6-well plate) to wet the entire membrane and kept in the incubator overnight to let cells adhere to the membrane. The next day, media was added to each well to float membranes in wells. The membranes with membranes were maintained in the incubator to become confluent.

The microfluidic chip consisted of two PDMS compartments (the top channel as the apical side and the bottom channel as the basal side), two adhesive pieces that were cut and attached to the top, and the bottom channels, and a porous polyester membrane with Calu-3 cells. Figure 6A and Figure 6B show the fabrication steps of the top and the bottom channels and their attachment to the cut adhesives. A fully assembled device (Figure 6C) was filled with DI water contained dyes, as shown in Figure 6D, to ensure that there was no leakage after forming wet bonding between the PDMS compartments and the porous polyester membrane using wet bonding. In addition, to make sure that the wet bonding between the membrane and both channels would not lead to any leakage, the channels were perfused with fluorescent dyes and imaged as presented in Figure 6E and Figure 6F. Figure 6G shows the cells on a porous membrane after assembling the chip using bright-field microscopy. Moreover, the viability of Calu-3 cells one hour after the transfer was assessed using Calcein AM and DAPI (Figure 6G).

After transferring a membrane with adhered Calu-3 cells to the chip, both apical and basal channels were filled with media and maintained in an incubator until the cell became 100% confluent. Upon their confluency, the media from the apical side was removed, and the cells were introduced to the air-liquid interface (ALI). The cells were kept under ALI conditions before stimulating them with IL-1β. In the next step, the chips were basally perfused for 12 hours at a flow rate of 20 μL min^−1^ with media containing IL-1β (5ng/mL) and media was also added to the apical side with the same concentration of IL-1β (the control experiment was conducted without adding IL-1β to the media). Similar to single-channel chips, IL-8 and IL-6 as inflammatory cytokines were measured and compared to controls.

Similar to the single-channel chips, stimulation with IL-1β (5ng/mL) caused inflammatory cytokine responses in both apical and basal sides. IL-8 production on the apical side was increased by ~ 50 % compared to the control samples (Figure 7A). However, IL-1β (5ng/mL) stimulation caused a higher increase (~ 5-fold) in IL-6 production on the apical side (Figure 7B). In addition, the production rate of IL-8 and IL-6 on the basal side of the cells was also raised by ~ 5.4 and 2 times, respectively (Figure 7C and Figure 7D).

After IL-1β (5ng/mL) stimulation, the cells were stained using Calcein AM and DAPI to demonstrate that the cells remained viable, as shown in Figure 7E.

## 4 Discussion

Current research in the field of airway epithelial cell biology has seen recent advances with cell culture miniaturization to incorporate different cell techniques that improve throughput, cut costs of reagent use and provide researchers with more customization options [13]–[17]. In addition, significant progress has been made on microfluidic organ-on-a-chip devices that integrate forces into cell culture, which are absent when studying cells in conventional static culture systems[9], [16]. While these methods provide alternative options for researchers to study airway epithelial cells, they have limitations. Many miniaturization methods require technically difficult procedures and access to equipment and training that is not readily accessible to biomedical researchers[7], [17]. Furthermore, currently studied organ-on-a-chip models often require pre-seeding cells into a pre-defined structure cast out of a mold. These models offer limited flexibility in design and do not allow researchers easy access to cells after perfusion culture. Our data demonstrate that using commercially available adhesives has several applications to address the aforementioned limitations. We demonstrate a method of developing a microfluidic culture device using adhesives that do not rely on the pre-seeding of cells. The dynamic microfluidic model system was robust in a variety of experimental conditions and supported cell viability and functional airway epithelial immune responses.

Several commercially available adhesives were tested for applicability in cell culture and other applications. Although no significant cytotoxic effects were found between the four different tested adhesives (Figure 1), some limitations were discovered that led to the pursuit of using the acrylic-based ARclean® 90716 adhesive for model development. In testing, acrylic-based Scotch™ 7951 lost tactile properties following prolonged submersion in fluid, rendering it unusable for cellular growth in submerged media. Polyester-based ARclad® 7535-12 was functionally suitable for experimental use but displayed autofluorescence using the DAPI light cube while performing fluorescent microscopy imaging. Finally, silicone-based ARclean® 8932EE was relatively elastic, causing it to poorly retain the desired shape and also did not remain functionally tactile following prolonged submersion in media. ARclean® 90716 fulfilled the desired properties for applications in microfluidic device manufacturing, and because no cytotoxic differences were seen between the tested adhesives, it was utilized for all experiments.

Our proposed microfluidic model was user friendly to assemble and is made of materials that are applicable to microscopy and other imaging techniques (Figure 2). Based on the application, the design and size of the chip can be easily modified without impacting the fabrication process. The functional components of the model include adhesives, a transparent polyester membrane, and a PDMS lid that are all transparent with little interference in the wavelengths necessary for basic cellular imaging, with the exception of some adhesives. The model is also adhered to a glass microscopy slide, which, while being optically clear, also offers structural support to improve handling the chip and makes the chip easily insertable into the majority of microscope imaging platforms. PDMS has gas diffusion properties that allow cells to grow in the chip to be continuously exposed to environmental oxygen and carbon dioxide that are necessary for growth. Thus, the model has potential applications in studies monitoring real-time changes in cell morphology, migration, and gene/protein expression if used in a temperature and gas-controlled microscope. Moreover, we showed that a more advanced model with dual channels could be developed to expose airway epithelial cells to room air while being basally perfused (Figure 6). Such a model enabled us to assemble lung-on-a-chip devices for studying their immune responses to IL-1β stimulation (Figure 7).

ARclean® 90716 was able to maintain a fluid-tight seal following submersion in media for one week at flow rates from 10-250 μL/min (Figure 3A), which exceed the necessary flow rate for experimental purposes. Therefore, the model is capable of undergoing increased fluid flow rates to test extreme shear stress responses on cells. It is proposed that the variability in burst pressure seen as the chip is submerged for a greater number of days is due to media leaking in between the adhesive and its protective seal. For HAECs, the typical pressure experienced by interstitial fluid pressure is 8mmHg[22]. Therefore, although increased variation was seen in burst pressure for the extended submersion conditions, the burst pressure achieved is far greater than necessary to study effects on HAECs with interstitial fluid flow conditions.

A beneficial feature of the adhesive based chip is its ability to avoid seeding cells into a fully fabricated fluidic model, as required by many reported epithelial chip-based culture models in the current literature [9], [16]. This feature of the microfluidic chip allows the cells to be grown using conventional submerged cell culture methods. This is a conventional method that would be in use already in many labs studying HAECs. Thus, the model does not require additional cell culture methods, and the cells could be exposed to chemical stimuli while in the submerged culture, then the effects of the stimuli could be studied with perfusion culture afterward. The top seal of the top adhesive was left adhered to the chip during submerged culture. After growth to confluency, the chip was removed, and the top seal of the top adhesive was removed. We were able to contain cellular adherence to the 0.5cm targeted growth chamber reproducibly by using this technique (Figure S1 in the supplementary). The method used to achieve targeted growth was also tested with additional shapes and resolutions for other applications. In lung-on-a-chip format, both top and bottom adhesives were not exposed to media resulting in better sealing after transferring membranes to the chips (Supplementary Video). This model was robust enough to perfuse both apical and basal channels right away after assembling the chips (Supplementary Video).

Calu-3 cells were shown to remain viable after growing in the assembled chip for 24 hours of perfusion at a flow rate of 20 μL/min (Figure 4). Although the model remains tightly adhered during perfusion applications, the PDMS lid was easily removed after perfusion, allowing the cells to be stained again for imaging. For lung- on-a-chip format, only one of the top or bottom adhesives should be used to be able to disassemble the chips if needed. This feature of the chip increases possible applications of study after exposing cells to dynamic forces. The proposed model demonstrated the ability to measure the inflammatory response (*i.e.*, IL-6 and L-8 secretion) of airway epithelial cells after exposure to an inflammatory stimulus (IL-1β, Figure 5 and Figure 7). These results suggest that the model is valid for testing immune response in HAECs.

**Figure 5:**
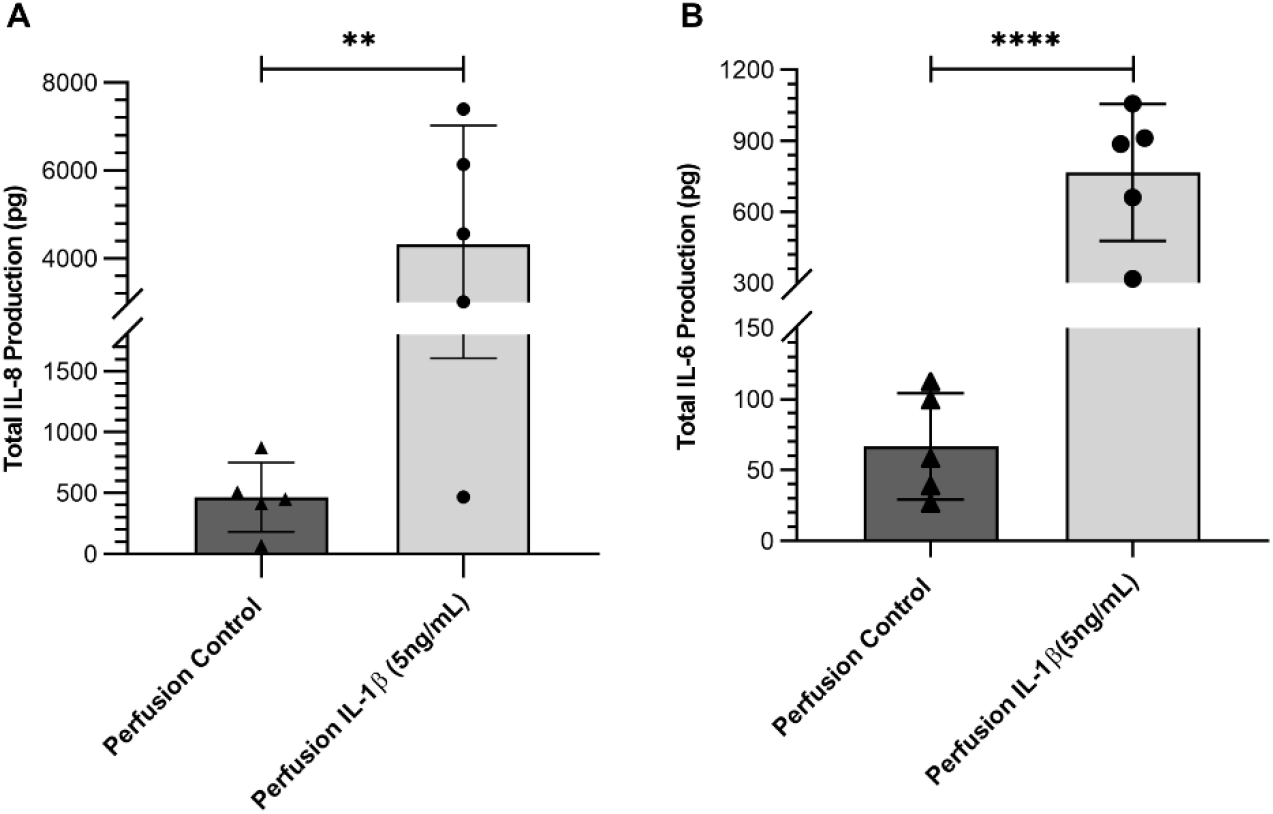
Comparison of Calu-3 IL-6 and IL-8 release after stimulation with IL-1β (5ng/mL) in a dynamic cell culture system. (A) IL-8 release expressed as total IL-8 (pg) calculated by adjusting for total exposure media volume. (B) Total IL-6 production. Results are displayed as mean ± SD. One-way ANOVAs were performed with a Tukey’s post hoc test for multiple comparisons. (P<0.01**, P<0.0001****). n=5

**Figure 6:**
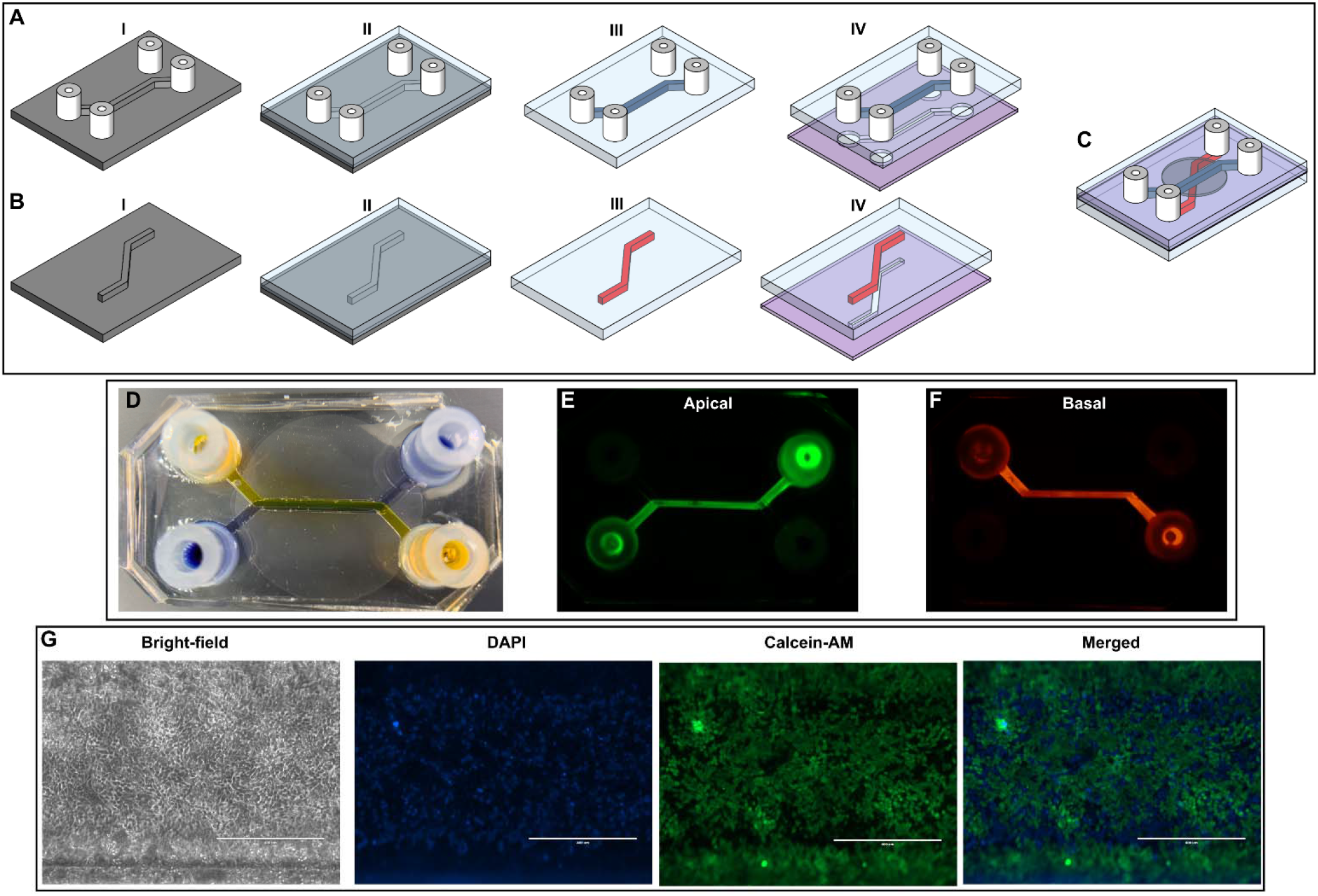
Assembly process of the microfluidic chip, a porous polyester membrane with Calu-3 cells, and the Viability assay of Calu-3 cells after assembling the chip. (A) the fabrication process of the top channel (apical side): (I) four 1-cm silicone tubes were placed onto the mold as inlets/outlet, (II) PDMS was poured on the mold and cured at 65 °C, (III) the cured PDMS was removed from the mold and inlets/outlets were to checked for any residual PDMS, (IV) an adhesive with the desired pattern similar to the design of the top channel was cut and attached to the top channel, (B) the fabrication process of the bottom channel (basal side): (I, II) PDMS was poured on the mold and cured at 65 °C, (III) the cured PDMS was peeled off from the mold, (IV) an adhesive with the desired pattern similar to the design of the bottom channel was cut and attached to the bottom channel. (C) the membrane was transferred onto the bottom channel and was sandwiched between two layers using adhesives. (D) the image of the chip (lung-on-a-chip) that was filled with dyes. (E, F) the apical and basal compartments were filled with a fluorescent dye and examined for any possible leakage after forming a wet bonding. (G) bright-field and fluorescent microscopies of Calu-3 cells after transferring a porous polyester membrane with cells to the chip. Scale bars are 400 μm.

**Figure 7:**
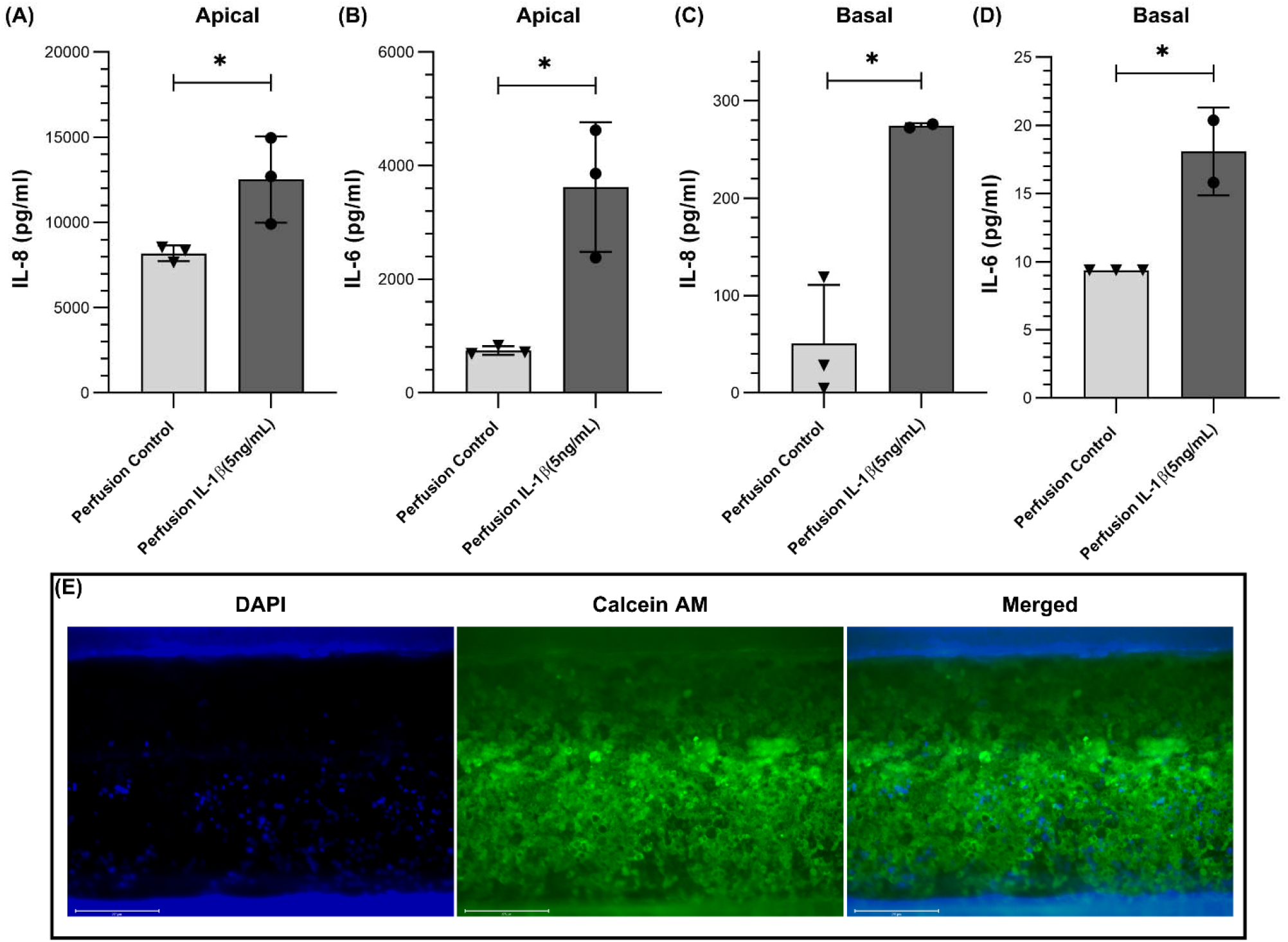
Comparison of Calu-3 IL-6 and IL-8 release after stimulation with IL-1β (5ng/mL) in a lung-on-a-chip format: (A) IL-8 production on the apical side. (B) IL-6 production on the apical side. (C) IL-8 production on the basal side. (D) IL-6 production on the basal side. (E) Qualitative Calu-3 cell viability after stimulation with IL-1β using Calcein AM and DAPI.

For future works, this fabrication technique can be used to integrate sensors in a micro-wire format, and the top part can be later attached to the bottom channel and a membrane (Figure S2 in the supplementary). After the assembly of the chips, the top channels will not be perfused with media to introduce the cells to ALI as well as protecting sensors from fouling. We used this fabrication process and fabricated such a device, as shown in Figure S3 in the supplementary. The assembled device was successfully perfused without any leakage.

## 5 Conclusion

In this paper, we have developed and tested an adhesive-based cell culture system that enabled us to pattern cells on a membrane using a patterned-cut adhesive as a mask. First, we showed that these adhesives were not toxic and did not compromise the cells’ morphology. We created various patterns on adhesives and could grow cells onto them. The cultured cells on these patterned adhesives were imaged after removing their top lining layers and transferring the desired patterns to the cells. This method was used to grow cells in a targeted area and was attached to a microfluidic chip for perfusion. Moreover, we advanced this method to fabricate lung-on-a-chip devices. The chips were perused and stimulated by IL-1β for 24 or 12 hours, showing a rise in IL-6 and IL-8 for both single-channel devices of lung-on-a-chip devices. The results suggest that this method can be used to grow cells outside of a microfluidic chip, which can be transferred later to a microfluidic channel for further analysis. Finally, we proposed that our technique can be utilized to fabricate lung-on-a-chip devices with integrated sensors so that the sensors will not be fouled as a result of not being exposed to the cell culture media.

## 6 Methods

### 6.1 Reagents

Calu-3 human adenocarcinoma airway epithelial cells (HTB-55™, ATCC®, USA) were cultured in alpha minimum essential medium (αMEM, Corning®, USA) supplemented with fetal bovine serum (10%, FBS, WISENT Inc., Canada) and antibiotic-antimycotic (100U/mL penicillin, 100μg/mL streptomycin, 0.25μg/mL amphotericin B, Gibco®, USA) and HEPES buffer (10 μM, Gibco®, USA). HBEC6-KT cells were cultured in Keratinocyte-SFM medium (KSFM, Gibco®, USA) supplemented with epidermal growth factor (EGF, 0.8ng/mL), bovine pituitary extract (BPE, 50μg/mL) and antibiotic-antimycotic (100U/mL penicillin, 100μg/mL streptomycin, VWR®, USA)[23]. Trypsin (0.05%, 0.53mM EDTA in HBSS) was obtained from Corning®, USA. Recombinant Human IL-1β was obtained from Peprotech, USA (#200-01B). The IL-6 and IL-8 enzyme-linked immunosorbent assay kits were obtained from R&D Systems, USA (#DY206 and #DY208, respectively). Trypan blue dye was obtained from ThermoFisher Scientific, USA. Lactate dehydrogenase kit was obtained from ThermoFisher, USA (#C20301). The adhesives used were ARclad® 7535-12, ARclean® 90716, ARclean® 8932EE (Adhesives Research Inc., USA) and Scotch™ 7951 (3M®, USA). All adhesives were sterilized with ethanol (70%) and ultraviolet radiation for 20 minutes before use. Porous polyester membranes (0.4μm, transparent, #PET0413100) were obtained from Sterlitech, USA. Polydimethylsiloxane (PDMS) was obtained from Dow Chemical Company, USA.

### 6.2 Microscopy and tools

All microscopic images were taken with the EVOS M7000 microscope using GFP, DAPI, and CY5 light cubes (ThermoFisher, USA). GFP, DAPI, and CY5 light cubes were used to visualize Calcein AM, Hoechst, and NucRed™ Live 647 ReadyProbes™ Reagent dyes, respectively. Calcein AM is a colorless dye that is converted to a green-fluorescent Calcein molecule after live cells’ cytosolic esterase enzymes cleave of the AM group. Therefore, it only stains live cells green-fluorescent. Hoechst and NucRed™ Live 647 ReadyProbes™ Reagent both stain the nuclei blue-fluorescent or red-fluorescent, respectively, of dead or alive cells.

Varying shapes of growth area or adhesives were designed in Silhouette Studio V4 software and cut out of ARclean® 90716 with a Silhouette CAMEO® 3.

### 6.3 Cell Culture and Viability Analysis

Calu-3 and HBEC6-KT cells were cultured in an incubator at 37°C with 5% carbon dioxide air on a two-day feeding cycle. For perfusion experiments, Calu-3 cells were cultured under submerged monolayer conditions on adhesive assembled chips with the top seal of the adhesive remaining on. The top seal was removed after confluency was achieved, and the PDMS lid was adhered to the chip. The cells were perfused with media at a flow rate of 20μL/min for 24 hours. The entire microfluidic device was incubated at 37°C for the duration of perfusion using a benchtop water bath.

Viability quantitative analysis was performed on Calu-3 and HBEC6-KT cells using Calcein AM and Hoechst dyes. After imaging the cells, the PBS solution was removed, and 500μL of trypsin was placed on the cells and incubated for 10 minutes at 37°C. The resuspended cells were centrifuged at 1,200 x g for 6 minutes and were resuspended with trypan blue for live/dead analysis. In addition, an LDH assay was performed to assess cell viability following the manufacturer’s directions. As a positive control, a control well of Calu-3 or HBEC6-KT cells were lysed to release all cytoplasmic LDH.

To quantify targeted Cellular Growth on a porous membrane, adhesive chips were assembled with ARclean® 90716 with the top seal remaining on. Calu-3 cells were grown to confluency and were incubated with Calcein AM at 37°C for 20 minutes. Images of the cells were taken before and after adhesive seal removal.

Calu-3 cells were grown in chips and incubated with Calcein AM and NucRed™ Live 647 ReadyProbes™ Reagent at 37°C for 20 minutes to assess the cells’ viability under perfusion. After imaging, a PDMS top was affixed, perfusion initiated (20μL/min), followed 24h later by imaging.

### 6.4 Flow Rate Validation and Burst Pressure Measurement of Microfluidic Model

Chips were submerged in αMEM 37°C for 1, 3, or 7 days. Following submersion, the top adhesive seal was removed, and the PDMS lid adhered. The chips were connected to a syringe pump with a pressure transducer that included monitoring system pressure over time (USB Output Pressure Transducer, Cat# PXM409-USBH, OMEGA Engineering Inc., USA). A syringe pump was set to 500μL/min and run in the chip until burst pressure was achieved. The final burst pressure was recorded.

To ensure the reversible bonding between the PDMS lid and the substrate was robust during the experiments, the burst pressure was evaluated, and the outlet flow was measured to show that there was no leakage under the operating conditions. Chips were submerged in αMEM at 37°C for one week. Following submersion, chips were removed from the media, the top adhesive seal was removed, and the PDMS lid was adhered to. Chips were connected to a syringe infusion pump (Harvard Apparatus, USA), with water run through, effluent collected, and mass measured. Chips were subjected to rates of 10, 50, and 250 μL/min for 30 minutes at each rate. Effluent mass was recorded at 5-minute intervals for a measure of volume.

### 6.5 IL-1β Stimulation and ELISA Analysis

Calu-3 cells grown in a static 96-well culture plate were grown with either control or media containing IL-1β (5ng/mL) for 24 hours. Calu-3 cells grown in the adhesive assembled perfusion chip were perfused control media or media containing IL-1β (5ng/mL) for 24 hours at a flow rate of 20μL/min. The IL-6 or IL-8 release was converted to total IL-6 or IL-8 (pg) to normalize for volume differences between static and dynamic conditions. For static culture conditions, the total collected volume was 200μL. For perfusion culture conditions, the total collected volume was 28.8mL (20μL/min*60min*24hr). The total cytokine released was calculated by applying the following formula:

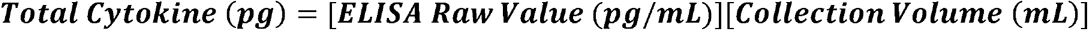

