## Supplemental Material for "An Adhesive-Based Fabrication Technique for Culture of Lung Airway Epithelial Cells with Applications in Microfluidics and Lung-on-a-Chip"

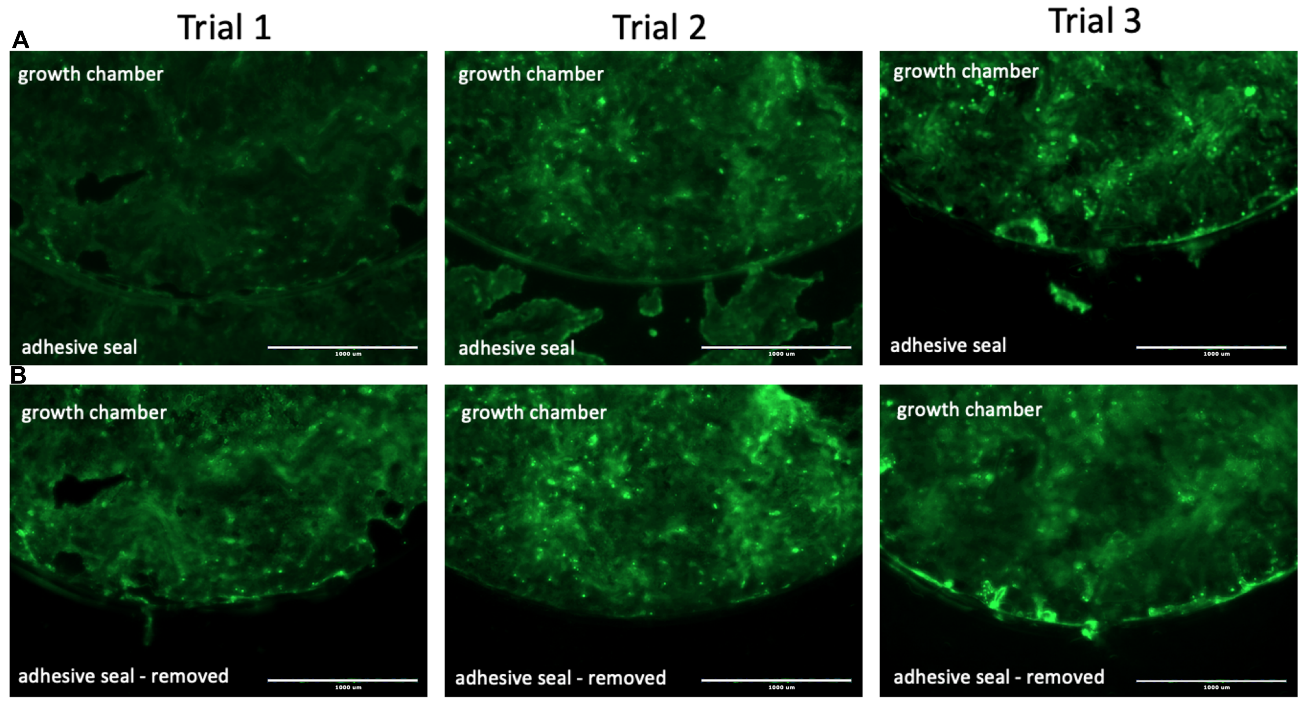

Figure S1: Maintenance of Calu-3 cellular growth in the target growth chamber following adhesive seal removal. Adhesive assembled chips were made with ARclean® 90716 with a 0.5cm diameter growth chamber. Calu-3 cells were seeded into the culture dish and grown to confluency. The cells were incubated with calcein AM and imaged with a GFP light cube prior to removing the adhesive seal and after the adhesive seal was removed. Images were taken at 40X magnification, scale bars represent 1000 µm.

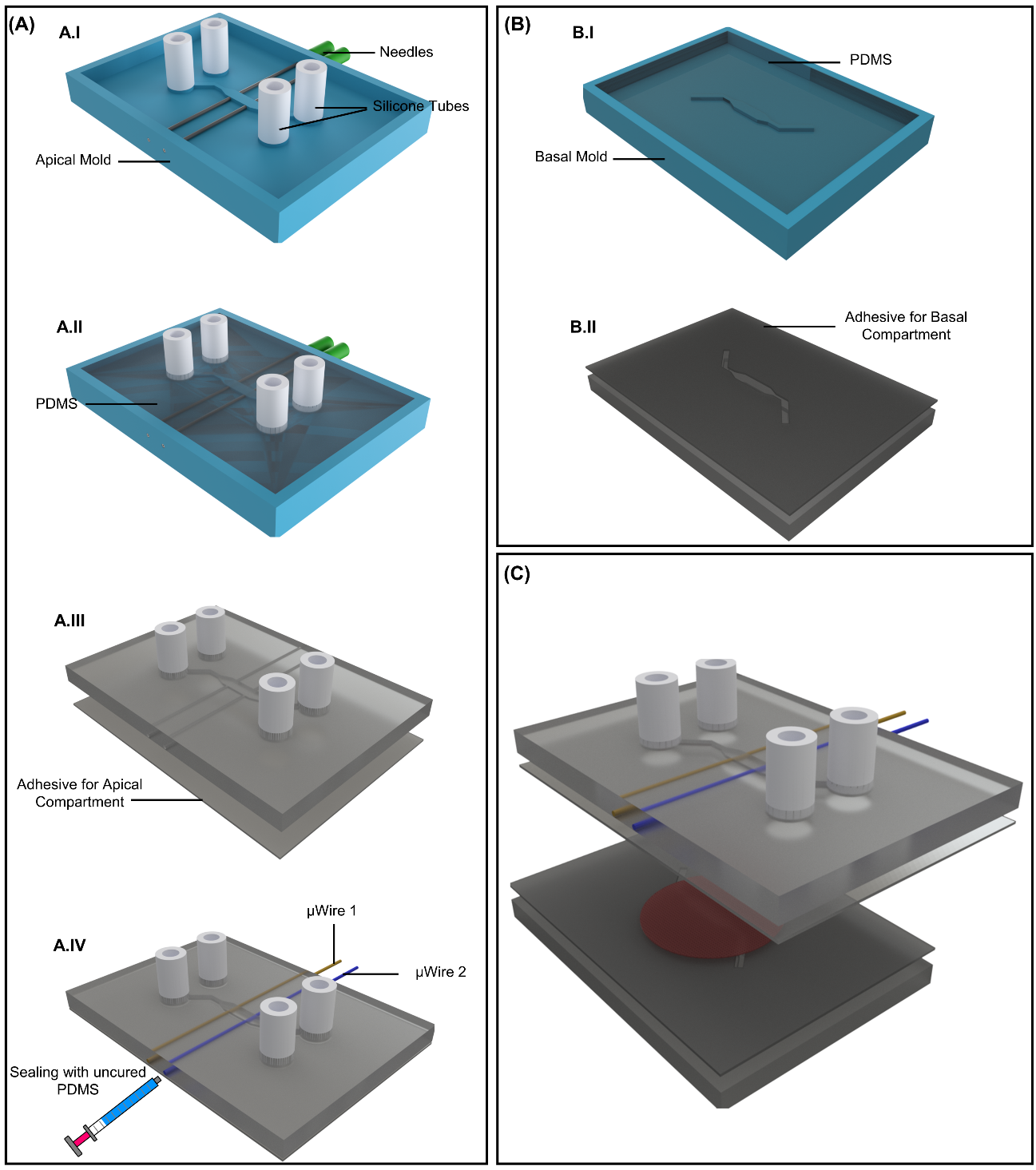

Figure S2: A proposed fabrication process for lung-on-a-chip devices with integrated sensors in wire format: (A) the fabrication process for the apical side with the integrated micro-wires. (B) the fabrication process for the basal compartment. (C) the unassembled device.

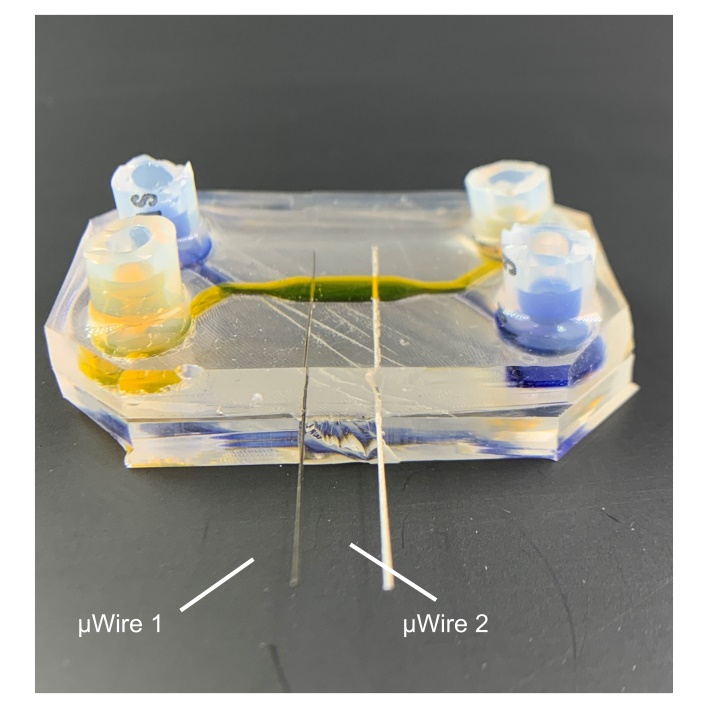

Figure S3: A fully assembled lung-on-a-chip device with integrated micro-wires using the adhesive fabrication technique proposed in this work. Both apical and basal channels were filled with dyed water.

1. Both authors contribute equally to this work.
